# Elevation of Gut-Derived p-Cresol During Spaceflight and its Effect on Drug Metabolism and Performance in Astronauts

**DOI:** 10.1101/2020.11.10.374645

**Authors:** Michael A. Schmidt, Cem Meydan, Caleb M. Schmidt, Ebrahim Afshinnekoo, Christopher E. Mason

**Affiliations:** Advanced Pattern Analysis & Countermeasures Group, Boulder, CO USA; Sovaris Aerospace, Boulder, CO USA; Department of Systems Engineering, Colorado State University, Fort Collins, CO USA; Department of Physiology and Biophysics, Weill Cornell Medicine, New York, NY, USA; The HRH Prince Alwaleed Bin Talal Bin Abdulaziz Alsaud Institute for Computational Biomedicine, Weill Cornell Medicine, New York, NY, USA; The WorldQuant Initiative for Quantitative Prediction, Weill Cornell Medicine, New York, NY, USA; The Feil Family Brain and Mind Research Institute, Weill Cornell Medicine, New York, NY, USA

**Keywords:** NASA, Spaceflight, *p*-Cresol, CYP4502E1, Metagenome, International Space Station, NASA Twins Study, Multi-Scale Omics, Precision Medicine

## Abstract

Metabolites produced by enteric microbes may have important effects on astronauts on long-duration missions. The NASA Twins Study provided the most comprehensive multi-scale omics data to date from which to extract molecular features of potential clinical significance to spaceflight. From the multivariate data, we identified an elevation of the uremic toxin *p*-cresol, which is produced by gut microbial fermentation of dietary tyrosine. *p*-Cresol has adverse metabolic effects via depletion of the hepatic sulfur pool, which impacts metabolism of drugs, endogenous metabolites, and xenobiotics. Moreover, *p*-cresol reshapes gut microbial community structure by facilitating survival of species such as clostridia and inhibition of butyrate producers. Spaceflight may also impact the genes responsible for the metabolism of *p*-cresol (e.g. SULT) and for the safe metabolism of common drugs used in space, such as acetaminophen (e.g. SULT, CYP4502E1, GST). Understanding *p*-cresol production and its related molecular networks in astronauts may lead to precision medicine advances that enhance astronaut safety and performance on long-duration missions.

## 2. Introduction

The human intestinal microbiota is well known for its complex effects on host metabolic phenotypes. However, the dynamics of human-microbial co-metabolism in long-duration human spaceflight is not well understood. The NASA Twins Study represents the longest study of a human in space that incorporates longitudinal measures derived from multi-scale omics. The study incorporated over 300 samples collected across 19 time points before, during and after flight. Data were generated using multiple modalities to examine human and microbial biology, including isolation of saliva, skin, plasma, stool, urine, peripheral blood mononuclear cells (PBMCs), and immune cells.

*p*-Cresol is of specific importance to spaceflight because of its broad mechanistic dynamics. Once *p*-cresol produced in the gut, it is absorbed across the mucosal barrier, enters the liver, and is metabolized via sulfation or glucuronidation. This leads to the formation of *p*-cresol sulfate and *p*-cresol glucuronide. Persistent elevation of *p-*cresol sulfate over time can deplete the hepatic sulfur pool, and rob it of the sulfur groups needed to metabolize drugs (acetaminophen, levothyroxine, amoxicillin, morphine, hydrocortisone, etc.), endogenous compounds (estrogens, androgens, catecholamines, bile acids, thyroid hormones, etc.), and xenobiotics.

Using acetaminophen as an example for altered drug metabolism, a depleted sulfur pool is problematic because inadequate sulfation of acetaminophen can result in formation of the hepatotoxin *N*-acetyl-p-benzoquinone imine (NAPQI). Unconjugated NAPQI binds to proteins and subcellular structures, and induces rapid cell death and necrosis that can lead to liver failure. Ordinarily, adequate supply of glutathione (GSH) permits formation of an NAPQI-GSH conjugate for safe excretion. However, *p*-cresol has been shown to lower GSH, which suggests the presence of *p*-cresol may render the formation of NAPQI more probable (Edamatsu et al., 2018). (Figure 1).

**Figure 1.**
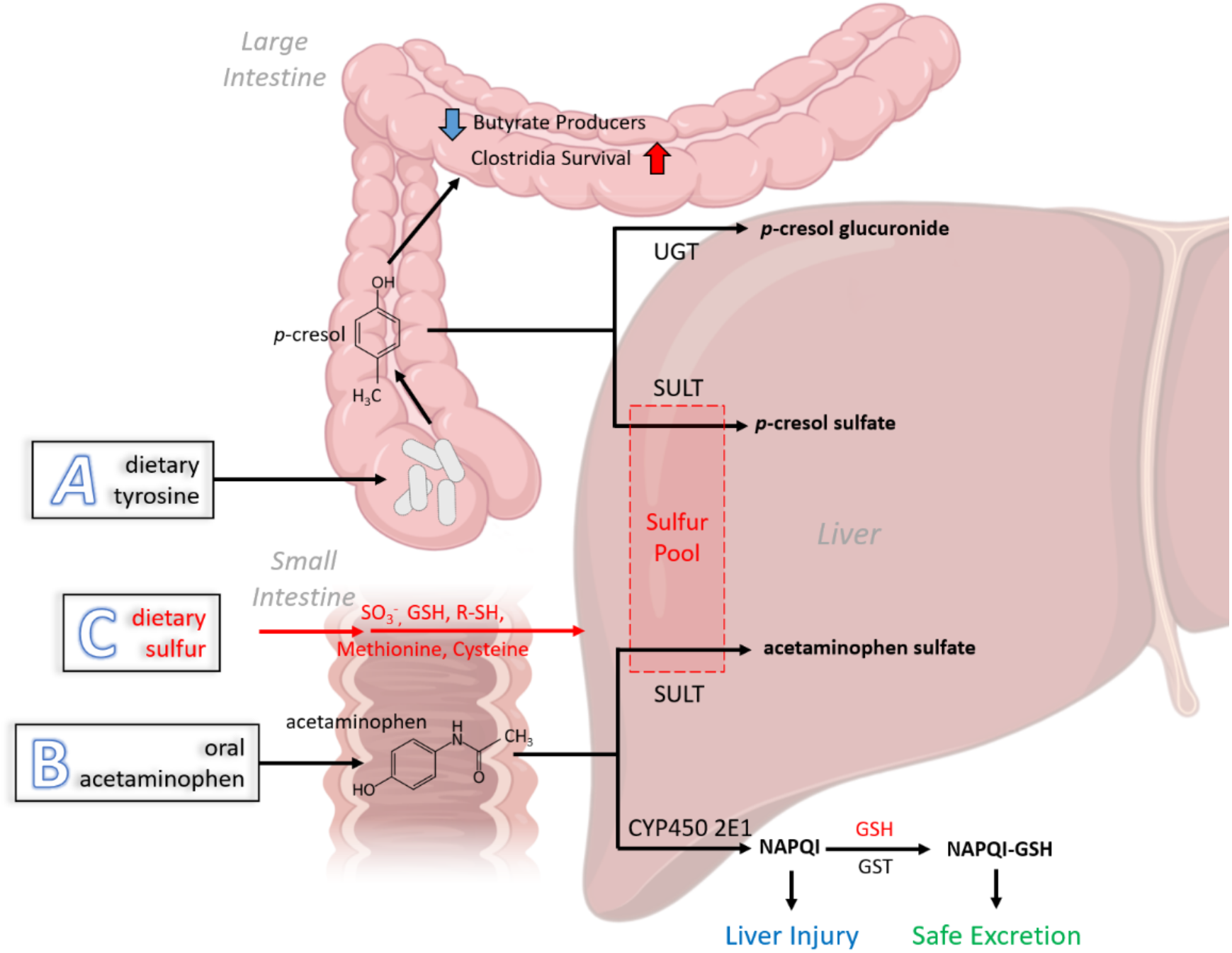
*p*-Cresol Formation and its Impact on Molecular Networks in Astronauts. The figure shows three processes with intersecting relationships: A) Dietary tyrosine is converted by colonic microbiota to *p*-cresol. *p*-Cresol is absorbed into hepatic circulation, where it is metabolized by UGT into *p*-cresol glucuronide and by SULT into *p*-cresol sulfate. *p*-Cresol sulfation utilizes and requires a sufficient hepatic sulfur pool. *p*-Cresol that remains in the colon inhibits butyrate producing microbes, while facilitating survival of clostridial species (*p*-cresol producers); B) oral acetaminophen is absorbed through the small intestine and transported to the liver where it is metabolized via SULT into acetaminophen sulfate (Note: the acetaminophen glucuronidation path is omitted for simplicity). This process directly competes with *p*-cresol for SULT activity and for sulfur molecules. Up to 15% of acetaminophen is also metabolized by CYP4502E1 into NAPQI. In the presence of adequate glutathione and active GST, NAPQI is rendered harmless and excreted. In the absence of adequate glutathione, NAPQI is hepatotoxic; C) Dietary sulfur groups (sulfur pool) are provided in the form of SO3-, glutathione, cysteine, methionine, and others. These replenish the hepatic sulfur pool needed to remove drugs, endogenous compounds, and xenobiotics. The presence of *p*-cresol can deplete the hepatic sulfur pool needed for the safe metabolism of drugs like acetaminophen in space. UGT: UDP-glucuronosyltransferase; GSH-glutathione; SULT-sulfotransferase; NAPQI-*N*-acetyl-*p*-benzoquinone imine; RSH-generic sulfhydryl compound; GST-glutathione-*S*-transferase.

**Figure 2.**
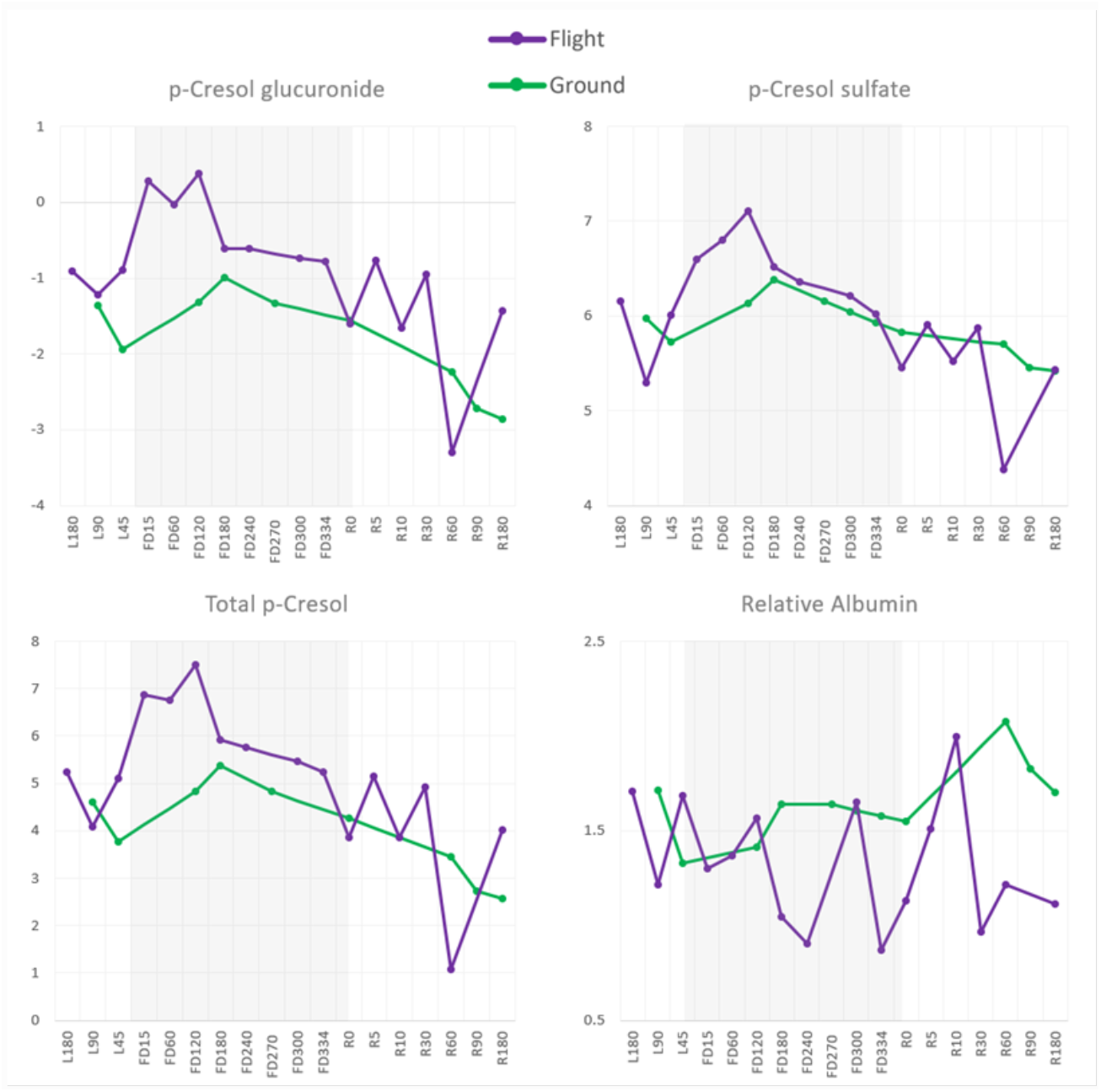
Increase in Plasma *p*-Cresol During One Year in Space. Ground control (green) levels for p-Cresol are shown relative to the flight subject (purple) across days before launch (L-), during the flight days (FD), and days post return to Earth (R).

It is important to note that hepatic cytochrome P450 2E1 (CYP450 2E1) is responsible for roughly 15% of the metabolism of acetaminophen and is the enzyme responsible for NAPQI formation. The process of phase II sulfur conjugation of acetaminophen is governed by sulfotransferases (SULT), while NAPQI conjugation is accomplished via glutathione-*S*-transferases (GST). Thus, we also examined CYP450 2E1, SULT, and GST RNA transcripts during one year in space. Finally, *p*-cresol impacts gut microbial diversity, favoring *p*-cresol-forming organisms and inhibiting butyrate-forming organisms.

*p*-Cresol is of general importance to spaceflight, since it has been associated with reactive oxygen species generation, endothelial dysfunction, epithelial-to-mesenchymal transition, leukocyte– endothelial interaction, deterioration of cardiac cell functional capacity, expression of tissue factor in vascular smooth muscle cells, renal dysfunction, and insulin resistance (Vanholder et al., 2014). Moreover, *p*-cresol may have central nervous system effects, having recently been shown to be elevated in the cerebrospinal fluid in Parkinson’s disease compared with controls (Bartlomiegj et al., 2020) and mechanistically associated with altered dopamine production (Pascucci et al., 2020). In a recent profile of 700 metabolites and concurrent assessment of global cognitive function, *p*-cresol was among the top three compounds differentiating the low from the high cognition group (Palacios et al., 2019).

For the reasons cited above, we assessed plasma *p*-cresol derived from gut microbial metabolism during one year in space, examined its associated molecular networks, and consider the implications for drug metabolism, astronaut health, and astronaut performance on long-duration spaceflight.

## Materials and Methods

### Sample collection

A pair of monozygotic twins were studied for 25 months, during which one subject (Flight subject, TW) spent 340 days aboard the International Space Station (ISS) while his identical twin (Ground subject, HR) remained on Earth. Subjects were male and aged 50 at the beginning of the study. Both subjects had different amounts of spaceflight exposure prior to the study (Flight subject=180 days total, Ground subject=54 days total). Multiple blood samples were collected (N_flight_=19, N_ground_=12) from both subjects beginning approximately 6 months prior to the launch date, during the 340 days aboard the ISS and 6 months after return.

All of the freshly collected samples aboard the ISS from the Flight subject were returned on the Soyuz capsule. Therefore, samples derived from ISS habitation were confounded by ambient return to Earth. To correct for the effects of ambient return on gene expression, blood samples were collected from an unrelated individual, which were then exposed to conditions that simulated the fresh and ambient return collections. The fresh and ambient samples from the control individual were processed and sequenced with the same protocol used for the study subjects.

### Data analysis

Data generated from the control individual was used to model and correct the effects of ambient return, using ComBat (Johnson et al., 2007; Leek et al., 2019) or multivariate analyses in differential expression, as previously described (Garrett-Bakelman et al., 2019). Blood sample collection was performed as previously described (Garrett-Bakelman et al. 2019).

## Results

### Plasma p-Cresol

Plasma *p*-cresol glucuronide and *p*-cresol sulfate increased in relation to baseline early into the spaceflight mission (TW) and began to decline mid-mission. Levels of *p*-cresol glucuronide increased in spaceflight in relation to the ground control condition. *p*-Cresol sulfate rose during spaceflight (TW) in relation to the ground control, though levels converged later in the mission.

## Discussion/Conclusion

This appears to be the first evidence that *p*-cresol sulfate and *p*-cresol glucuronide are elevated during spaceflight, based on untargeted analysis and measures of relative concentration. This may suggest changes in diet and/or gut microbial species that favors the conversion of dietary tyrosine to *p*-cresol. This may also reflect an unknown effect of spaceflight the converging factors that govern *p*-cresol production.

Should the elevation of *p*-cresol be found to be a repeatable phenomenon in long-duration spaceflight, as series of pre-mission or in-flight countermeasures warrant consideration. Countermeasure strategies to remediate elevated *p*-cresol in space that warrant consideration include 1) repletion of the sulfur pool, 2) dietary changes, 3) prebiotics, 4) probiotics, 5) synbiotics, and 6) pre-mission assessment of selected provisional biomarkers coupled with personalized intervention.

Attention to the sulfur pool should be among the lead considerations in addressing elevated *p*-cresol. *N*-acetylcysteine (NAC) is the accepted clinical means to treat acetaminophen poisoning in the emergency room setting, as it is known to reliably and rapidly replete the sulfur pool. While NAC is known to replenish the sulfur pool, it is also known to restore glutathione synthesis and protect the liver against NAPQI formation when acetaminophen is ingested (Ben-Shachar et al., 2012). Oral or intravenous glutathione can also be considered as a pre-mission countermeasure (glutathione being a tripeptide of cysteine, glycine, and glutamic acid). Alpha-lipoic acid has also been used to restore glutathione levels.

A general means to address *p*-cresol in space is to prevent its formation via dietary modification. Dietary restriction (tyrosine restriction) has been proposed as one means to effectively reduce *p*-cresol levels, as it reduces tyrosine intake from protein. However, the reduction in protein intake would have adverse consequences on muscle, while also further reducing the intake of sulfur-bearing amino acids, such as L-cysteine and L-methionine.

In one small study comparing vegetarians to omnivores consuming an unrestricted diet for one month, *p*-cresol sulfate excretion was 62% lower in vegetarians, which coincided with a fiber intake 69% higher and protein intake 25% lower than the *ad libitum* omnivores (Patel et al., 2012). Prebiotics are dietary substrates that distribute to the colon where they are acted upon by colonic microbes. These can take the form of high molecular weight polymers (soluble and insoluble fibers) that are consumed in gram amounts daily (e.g. resistant starches from potatoes) or polyphenolic compounds that are more commonly consumed in the hundreds of milligrams daily (e.g. proanthocyanidins from berries).

Resistant starch is a carbohydrate that resists digestion in the small intestine. Resistant starch has recently been shown over 8 weeks (20-25 g/day) to significantly lower *p*-cresol in humans (Khosroshahi et al., 2019). In Wistar rats on a tyrosine-rich diet, resistant starch was shown to lower urinary *p*-cresol levels to that of controls (Chen et al., 2016). In a 12-week feeding study, adult participants consumed 1) a control diet, 2) muffins with 10g/day pea hull fiber (4 weeks), and 3) muffins with 10g/day pea hull fiber and 15g/day inulin (6 weeks). Total fiber intake was 16.6 g/day, 26.5 g/day, and 34.5 g/day respectively. Providing pea hull fiber and inulin reduced serum *p*-cresol by 24% and by 37% in those with the greatest compliance (compliance being tied to greater inulin intake) (Salmean et al., 2015).

Probiotics are live microorganisms that are used to improve health, generally with the intention of restoring the gut microflora. Presently, there is little data on probiotics given alone, as a means to lower *p*-cresol in humans. In one study of 42 hemodialysis patients, participants were given 1.6 × 10^7^ CFU of *Lactobacillus rhamnosus* for 28 days (Eidi et al 2018). Mean serum *p*-cresol levels were lower after 4 weeks administration of *L. rhamnosus* when compared to the baseline (2.68 mg/dL versus 1.23 mg/dL; p=.034)

Synbiotics are merely a combination of prebiotics and probiotics. In one study of healthy volunteers, a synbiotic treatment, consisting of B. breve Yakult, L. casei Shirota and oligofructose-enriched inulin, reduced urinary *p*-cresol (a surrogate marker for colonic *p*-cresol) and promoted the growth of bifidobacteria (De Preter et al., 2007). In a study by Guida et al, participants with elevated *p*-cresol and kidney disease were provided a synbiotic consisting of lactobacillus and bifidobacterial species, plus fructooligosaccharides, inulin (6.6g/day), and tapioca. They showed a 40% reduction in *p*-cresol over 30 days (Guida et al., 2014).

Several limitations of this study warrant consideration. First, the small number of subjects and large number of variables (analytes) renders such high-dimensional data prone to overfitting. One method to address this limitation is through serial measures of the baseline, in-flight, and post-flight conditions, which was the approach used in this study. Second, the cohort of males is limiting on the generalizability of these results to females. Nonetheless, the matched cohort favors homogeneity, which is valuable in small N studies involving high-dimensional data. Third, the longitudinal measures of *p*-cresol derived from the untargeted metabolome were not fully quantitative.

Future directions consist of two paths. First, additional analysis of the existing NASA Twins data is currently underway, which is focused on key elements of the *p*-cresol molecular network. This consists of analysis of 1) Sulfotransferase RNA, 2) glutathione-*S*-transferase RNA, 3) serum albumin, 4) cytochrome P4502E1, 5) gut microbial genomic DNA and taxonomic shifts, 6) functional gene content of butyrate-producing gut microbial genes (urease, Uricase, tyrosine lyase, and hydroxyphenylacetate decarboxylase), and 7) functional gene content of *p*-cresol-producing gut microbial genes (phosphotransbutyrylase and butyrate kinase).

Second, future prospective studies in should be undertaken that attempt to further clarify the trajectory of *p*-cresol in space, along with its impact on the clinical and performance phenotype. This should include measures of *p*-cresol and associated measures that are fully quantitative. These associated measures would include but not be limited to 1) glutathione, 2) L-cysteine, and 3) L-methionine.

Finally, given the impact of *p-*cresol on the hepatic sulfur pool and the importance of a stable sulfur pool in space, the pre-mission analysis of these compounds may be warranted. Status of these measures may indicate whether active countermeasures are warranted today.

## Acknowledgements

We would like to thank the Epigenomics Core Facility at Weill Cornell Medicine, the Scientific Computing Unit (SCU), XSEDE Supercomputing Resources and funding from the Irma T. Hirschl and Monique Weill-Caulier Charitable Trusts, Bert L and N Kuggie Vallee Foundation, the WorldQuant Foundation, The Pershing Square Sohn Cancer Research Alliance, NASA (NNX14AH50G, NNX17AB26G), the National Institutes of Health (R01AI125416, R01MH117406, R01CA249054, R01AI151059, P01HD067244, P01CA214274), TRISH (NNX16AO69A:0107, NNX16AO69A:0061), the Bill and Melinda Gates Foundation (OPP1151054), the Leukemia and Lymphoma Society (LLS) grants (LLS 9238-16, Mak, LLS-MCL-982, Chen-Kiang). We would like to thank the NASA JSC Nutritional Biochemistry Lab personnel, especially YaVonne Bourbeau, who led efforts on this project for project coordination, sample collection, processing, and analysis; the NASA Human Research Program’s ISS Medical Projects element, Daniel Mollicone and Christopher Mott from Pulsar Informatics Inc., Emanuel Hermosillo and Sarah McGuire for support of the Cognition measures; Dr. Yael-Rosenberg Hasson from the Stanford Human Immune Monitoring Center for advice and support on cytokine profiling data; Keith Bettinger from Stanford University for support with the Stanford Twins data repository; Pawel R. Kiela DVM PhD, Daniel Laubitz PhD (University of Arizona), and Eunjoo Song (Northwestern University) for support of the Microbiome project sample collection; and Jiwan “John” Kim for laboratory support in targeted metabolomics. We thank Jason X.-Y. Yuan, MD, PhD at the University of Arizona, Tucson for providing his support and laboratory for the Tucson sample collections. Our thanks to Michael Snyder and Tejas Mishra of Stanford University for untargeted metabolomics and *p*-cresol analysis. We thank Julian C. Schmidt, M.S. (Sovaris Aerospace) for manuscript support. Finally, we thank Robert Hubbard, EdD, MA (Sovaris Aerospace) for manuscript review and illustrations. Elements of Figure 1 were created using BioRender.com.

## Statement of Ethics

The study participants have given their written informed consent. The study protocol has been approved by the research institute’s (both NASA and WCM) committee on human research, specifically IRB# 1309014347.

## Disclosure Statement

CEM is a cofounder and board member for Biotia, Inc., and Onegevity Health, Inc.

## Funding Sources

The study was supported by NASA: NNX14AH51G (All Twins PIs); TRISH: NNX16AO69A:0107 and NNX16AO69A:0061 (Mason); the Bert L and N Kuggie Vallee Foundation, the WorldQuant Foundation, The Pershing Square Sohn Cancer Research Alliance, and the Bill and Melinda Gates Foundation (OPP1151054) for funding (Mason).

## Author Contributions

M.A.S. and C.E.M. conceived the overall study objectives. M.A.S. and C.E.M. led the manuscript writing, organization, and editing efforts. M.A.S., C.E.M., C.M.S., and C.M. processed the data. C.M. and C.M.S. performed statistical analysis and multi-omics data integration. M.S. led molecular feature extraction, molecular target selection, pathway elucidation, and molecular network description. M.A.S., C.E.M., C.M.S., and C.M., E.A. performed the data interpretation, translation to biological meaning, summary of results, discussion, conclusion, and future directions.

## Notes

### Competing Interest Statement

The authors have declared no competing interest.

